# A high-throughput cell-based platform for Rhinovirus C research and antiviral drug discovery

**DOI:** 10.1101/2025.10.28.680218

**Authors:** Heyrhyoung Lyoo, Yeranddy A. Alpizar, Céline Sablon, Toon Röpke, Jasmine Paulissen, Madina Rasulova, Nathalie Thys, Chang-Soo Yun, Nam-Chul Cho, Kai Dallmeier, Pieter Leyssen, Soo-Bong Han, Johan Neyts, Hendrik Jan Thibaut

## Abstract

Rhinoviruses (RVs) are the most common cause of upper respiratory infections, yet RV-C is particularly associated with severe respiratory disease, including exacerbations of asthma and chronic obstructive pulmonary disease (COPD). Despite its clinical significance, RV-C research and antiviral development have been hindered by the lack of scalable and biologically relevant models for high-throughput screening (HTS). Here, we present the development, optimization, and validation of a scalable high-content imaging-based platform for RV-C research. Our approach combines engineered HeLa cells expressing CDHR3^C529Y^, a genetically stable mGreenLantern-reporter RV-C15 virus, and a phenotypic selection strategy to identify monoclonal cell lines with maximal and sustained permissiveness. This approach enabled assay optimization in both 96- and 384-well formats, achieving robust performance (Z′ > 0.7) and reproducibility across more than 100 runs with reference antiviral compounds. The platform was readily adapted to RV-A16, RV-B14, and additional RV-C types (C11 and C41), demonstrating broad applicability across rhinovirus species. A pilot screen of 10 240 compounds identified confirmed RV-C inhibitors, highlighting the readiness of the platform for integration into drug discovery pipelines. This platform provides a robust and scalable tool for systematic antiviral discovery and a foundation for mechanistic studies of RV-C replication.

**Significance statement:** Rhinovirus C (RV-C) is strongly associated with severe respiratory disease such as asthma and COPD exacerbations, yet its study has been limited by the absence of scalable cell-based systems. We developed a high-content imaging-based high-throughput screening platform that supports efficient RV-C replication in engineered CDHR3-expressing cells and enables systematic antiviral discovery. The platform integrates a genetically stable fluorescent reporter virus with phenotypically selected permissive monoclonal cell lines, achieving high reproducibility and scalability in both 96- and 384-well formats. Its adaptability across multiple rhinovirus species establishes a unified framework for antiviral testing and mechanistic investigation. A pilot screen of 10K compounds confirmed the applicability of the platform to drug discovery campaigns.

## Introduction

Rhinoviruses (RVs) are the most common cause of upper respiratory tract infections worldwide, yet their clinical burden extends far beyond the common cold. RVs contribute to bronchitis, pneumonia, exacerbations of asthma and chronic obstructive pulmonary disease (COPD), imposing a substantial global health burden. The discovery of RV-C species in the mid-2000s through molecular and phylogenetic analysis in pediatric cohorts with acute and severe respiratory illness revealed a previously unrecognized branch of rhinoviruses strongly associated with clinically significant disease (1–6). Despite its clinical importance, research on RV-C has lagged behind other RV species due to the lack of robust experimental systems.

Standard HeLa cell lines, which readily support RV-A and RV-B replication, fail to propagate RV-C (7), limiting both mechanistic studies and the development of antiviral strategies. Subgenomic replicon systems developed for several strains of RV-C allowed compound screening (8), but these systems do not recapitulate the full viral life cycle. Differentiated human airway epithelial (HAE) cultures later provided a major breakthrough, as they support productive RV-C infection (9, 10). However, these models are labor-intensive and costly, limiting their use for high-throughput screening. More recently, respiratory organoid models have been reported to support RV-C replication (11, 12), yet their complexity and high cost likewise limit scalability for drug screening. While these advanced models have deepened our understanding of RV-C replication, pathogenesis, and virus-host interactions (12, 13), none of them are suitable for systematic high-throughput antiviral discovery.

The identification of cadherin-related family member 3 (CDHR3) as the cellular receptor for RV-C (14) and its genetic association with asthma susceptibility (15) provided a breakthrough. The Tyr529 allele was shown to enhance receptor surface expression and confer increased susceptibility to RV-C infection (14, 16–19). HeLa cells engineered to express CDHR3-Tyr529 support RV-C propagation (14, 17), yet infection efficiency has remained insufficient for robust high-throughput screening (HTS) assay development.

High-content imaging (HCI)-based screening platforms have been successfully established for other respiratory viruses, including influenza A, RV-A16 (20), and severe acute respiratory syndrome coronavirus 2 (SARS-CoV-2) (21). These platforms typically rely on infection-based readouts (% infected cells) and require high infection efficiency to achieve stringent assay quality metrics, such as Z′ factor above 0.7. However, no such platform has been established for RV-C, leaving a critical gap in both mechanistic studies and drug discovery.

Here, we address this need by developing and validating a scalable HCI-based HTS platform for RV-C using engineered CDHR3^C539Y^-expressing HeLa cells and an mGreenLantern (mGL)-reporter RV-C15 virus. The reporter virus carries two adaptive mutations (VP1-T125K and 3A-E41K) that enhance heparan sulfate (HS)-mediated binding and replication in HeLa cells (22). Unlike previous approaches that prioritized receptor surface expression, we implemented a phenotypic selection strategy to identify monoclonal cell lines with stable and maximal permissiveness to RV-C15 infection. Using the selected monoclonal cell line, antiviral assay conditions in 96-well format were optimized to achieve robust Z′ values (>0.75), and subsequently miniaturized to a 384-well format to gain higher throughput. Both formats were validated with reference compounds across more than 100 assay plates, demonstrating robustness and reproducibility. The platform was further adapted to RV-A and RV-B species, as well as to additional RV-C types, including RV-C11 and RV-C41. Finally, a pilot screen of 10 240 compounds identified five confirmed RV-C inhibitors, demonstrating the readiness of the platform for integration into drug discovery pipelines. Together, this high-throughput cell-based platform provides a robust and biologically relevant tool that enables systematic antiviral discovery and mechanistic studies of RV-C replication.

## Results

### Generation of CDHR3-Expressing HeLa Cells and RV-C15a-mGL Reporter Virus

To date, no RV-C assay recapitulating the entire replication cycle has been available in a format suitable for high-throughput applications. To address this limitation, we generated a polyclonal HeLa cell line stably expressing the CDHR3^C529Y^ variant via lentiviral transduction. We specifically aimed to establish a biologically relevant screening platform that reflects viral replication dynamics rather than relying solely on receptor abundance. The detailed selection and characterization of highly permissive monoclonal lines are described in the following section (Fig. 1a).

**Figure 1.**
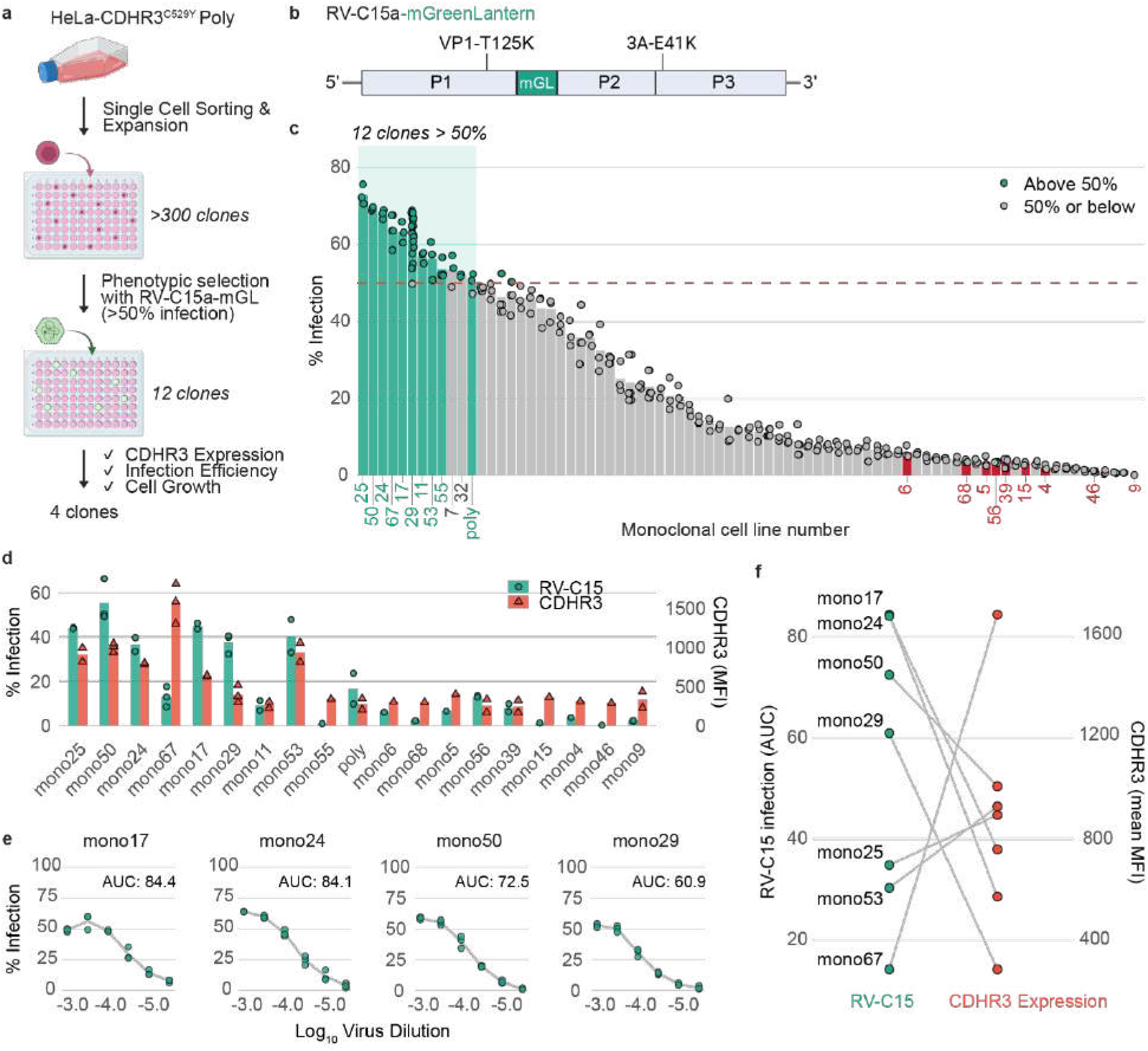
Generation of a stable RV-C15 reporter virus and phenotypic screening of permissive clones. (a) Schematic of HeLa-CDHR3^C529Y^ generation and workflow of phenotypic clone screening. Created with BioRender. (b) Schematic of RV-C15a-mGL virus genome orientation. (c) Primary screening of 80 monoclonal clones infected with RV-C15a-mGL. Infection efficiency was quantified at 3 dpi by HCI as the percentage of mGL-positive cells. Red dashed line indicates 50% infection threshold. Each circle represents an individual replicate (minimum n=3). Replicates with infection rates > 50% are highlighted in green circle, and replicates with ≤ 50% infection are shown in grey circle. Selected clones for further evaluation are indicated by colored bars and labels (green = permissive, red = negative). (d) Flow cytometry analysis of surface CDHR3 expression in permissive and non-permissive clones. Infection efficiency is shown as green circles (% mGL-positive cells), and CDHR3 expression as red triangles (mean fluorescence intensity; MFI). Each point represents the mean of two technical replicates from an independent experiment. Colored bars indicate the overall mean from all experiments (n=1 – 3). (e) Infection profiles of four stably permissive clones quantified as area under the curve (AUC) from dose-response infections with RV-C15a-mGL. Each circle represents an individual replicate (n=3). (f) Correlation between infection permissiveness (AUC) and CDHR3 expression (MFI) across monoclonal clones.

In parallel, we synthesized an improved version of the previously described RV-C15-GFP reporter virus (14), replacing GFP with mGreenLantern (mGL), a brighter and more photostable fluorescent protein (23) to enhance detection sensitivity in high-content imaging. Two adaptive mutations, VP1-T125K and 3A-E41K, previously reported to enhance heparan sulfate (HS) binding and replication in HeLa cells (22), were also introduced into the viral genome (Fig. 1b).

Polyclonal HeLa-CDHR3^C529Y^ cells were transfected with *in vitro*-transcribed RV-C15a-mGL RNA to produce viral stocks. The reporter virus replicated efficiently, and mGL fluorescence correlated with active replication site, as confirmed by anti-dsRNA staining (Fig. S1a). RT-PCR fingerprinting on viral RNA across passages demonstrated that the mGL insert remained genetically stable for up to at least two passages, which is sufficient for use in high-throughput assays (Fig. S1b). To confirm that the two adaptive mutations did not hinder natural infection characteristics, RV-C15a virus without mGL was also produced in the same polyclonal cells and tested in both human bronchial epithelial (HBE) and human nasal epithelial (HNE) cells grown at air-liquid interface, two systems previously shown to support RV-C replication (9), confirming the production of fully infectious virus particles (Fig. S1c). Replication of the virus was efficiently suppressed by the 3C protease inhibitor Rupintrivir (Fig. S1d), a well-characterized peptidomimetic inhibitor of human rhinoviruses (24). This validates the physiological relevance of the generated virus for downstream studies.

### Phenotypic Screening and Characterization of Highly Permissive Monoclonal Cell Lines

In contrast to the traditional selection approach based on high expression level, we applied a phenotypic screening strategy using the RV-C15a-mGL reporter virus to identify the most permissive cell clones (Fig. 1a). For this, more than 300 monoclonal cell lines were established by single-cell sorting from the polyclonal HeLa-CDHR3^C529Y^ population, of which 80 clones that exhibited robust growth after selection were chosen for initial screening. These clones were infected at a fixed dose of RV-C15a-mGL, and infection efficiency was quantified by high-content imaging (HCI) as the percentage of mGL-positive cells at three days post infection (dpi). Clones reaching >50% infection in at least two of three replicates were classified as highly permissive (Fig. 1c). Of those 11 clones that met this criterion, two (mono7 and mono32) were discarded due to slow growth.

To evaluate whether receptor abundance correlated with viral permissiveness, nine highly permissive clones (indicated in green in Fig. 1c), the polyclonal population, and eight non-permissive controls (indicated in red in Fig. 1c) were analyzed by flow cytometry across three independent rounds. Surface CDHR3 and HS expression was compared to infection efficiency, with mGL-positive cells serving as a proxy for virus infection (Fig. 1d, S2, and S3a). Several clones, including the polyclonal cell line, showed reduced growth or loss of permissiveness over time, consistent with possible clonal instability or phenotypic drift in culture. Of the nine permissive candidates, six (mono25, mono50, mono24, mono17, mono29, and mono53) consistently supported 40-60% infection. Interestingly, mono67 lost its permissiveness despite having the highest CDHR3 surface expression. Correlation analysis revealed only a moderate positive association between CDHR3 abundance (measured as median fluorescence intensity, MFI) and infection efficiency (Spearman’s ρ = 0.48, *p* = 0.036; Fig. S3b). Interestingly, there was a moderate negative correlation between HS abundance and infection efficiency (Spearman’s ρ =-0.66, *p* = 0.0028; Fig. S3b). Eight permissive candidates and three non-permissive controls were also screened against another C species reporter virus, RV-C11a-mGL, and the correlation analysis was performed to confirm the relationship between CDHR3 or HS abundance and RV-C11 permissiveness (Fig. S3d and S3e). As RV-C11a-mGL virus does not contain VP1 mutation that confers stronger binding to HS, no significant correlation between HS abundance and infection efficiency was observed (Spearman’s ρ =-0.47, *p* = 0.13), while a strong positive association with CDHR3 expression was found (Spearman’s ρ =0.81, *p* = 0.0016).

To confirm long-term stability and infection phenotype, seven clones were further tested in dose–response infections with RV-C15a-mGL in a 96-well format by HCI, and infection efficiency was quantified as the area under the curve (AUC) (Fig. 1e and S3f). After at least 10 passages, three clones (mono25, mono53, and mono67) showed a marked decline in permissiveness, with peak infection efficiencies below 30% (Fig. S3f), despite sustained moderate to high expression of CDHR3 in the cell surface (Fig. 1f). In contrast, four clones (mono17, mono24, mono50, and mono29) consistently supported >50% infection (Fig. 1e, 1f, and S3c) across 10 passages. This loss of permissiveness in several initially permissive clones emphasizes that receptor surface abundance alone does not guarantee sustained susceptibility to infection (Fig. 1f). Among the candidates, mono29 demonstrated the most stable cell growth and maintained its permissive phenotype for more than 30 passages (Fig. S3g), and was therefore selected as the lead cell line for further assay development.

### Development and Validation of a 96-Well Antiviral Assay

To determine assay conditions that balance high infection levels with sufficient cell viability, we tested a range of multiplicities of infection (MOI) and infection kinetics (Fig. 2a and S4a). Peak infection was reached at 72 hours post infection (hpi, Fig. 2a), corresponding with the highest amount of virus-induced cytopathic effect (Fig. S4a). All tested conditions yielded Z′ > 0.7, confirming the robustness of the assay for HTS applications (Fig. S4b). Based on these results, an MOI of 0.002 with a readout at 3 dpi was selected for subsequent antiviral assays, as this condition provided optimal infection efficiency, sufficient cell viability, and a high Z′ factor.

**Figure 2.**
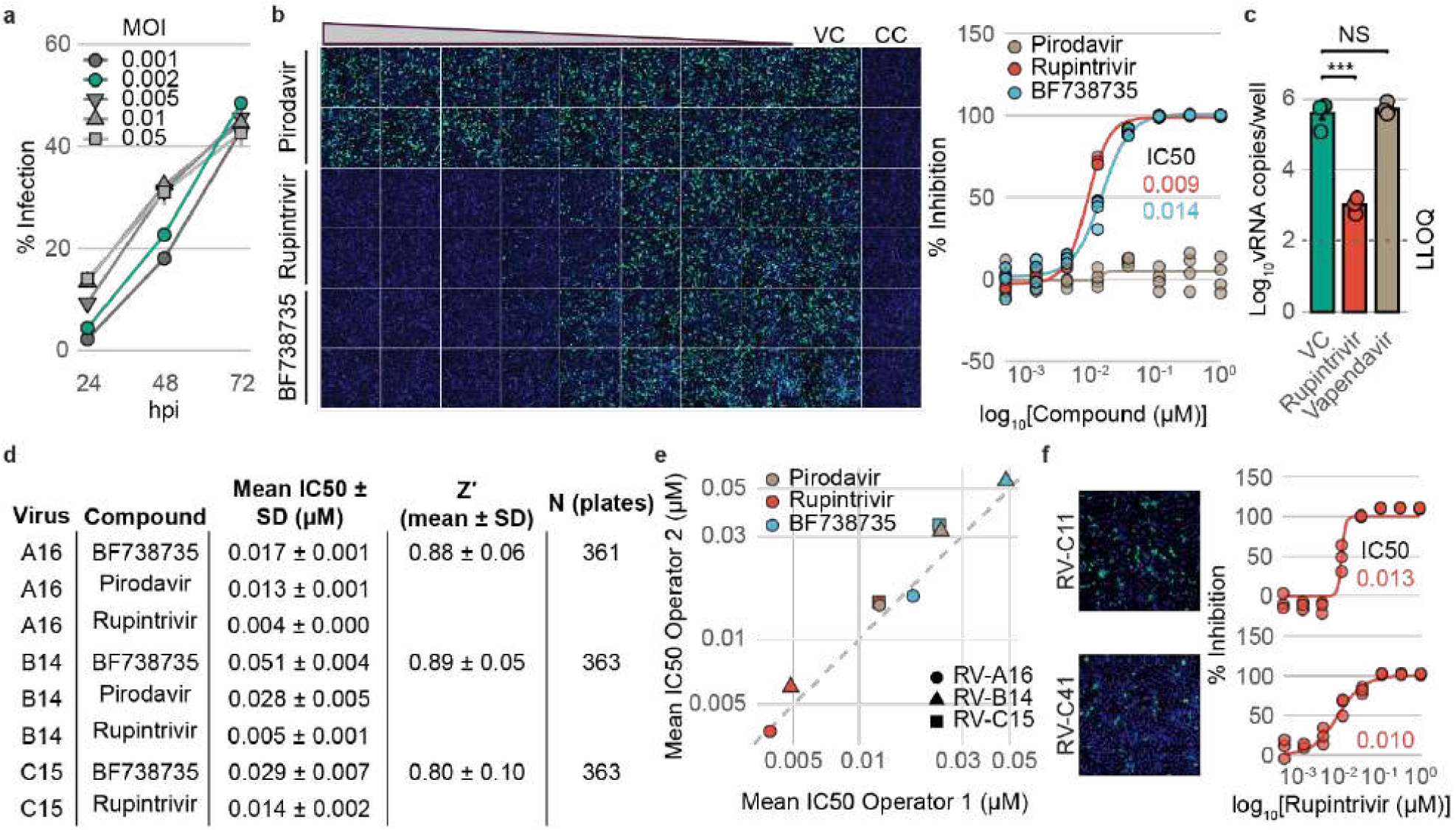
Development and validation of a 96-well RV-C antiviral assay. (a) Infection kinetics of RV-C15a-mGL in HeLa-CDHR3^C529Y^ mono29 cells at different MOIs and time points. Data are presented as mean ± SD (n=3). Selected MOI for antiviral assay is highlighted in green. (b) Pilot RV-C15a-mGL antiviral assay with reference compounds BF738735, Pirodavir, and Rupintrivir. Compounds were tested in 8-point dose-response assays starting at 1 µM. Assay plates were fixed at 3 dpi and analyzed by HCI. One representative plate is shown. Each point on the inhibition curves represents an individual replicate. Two operators performed the assay independently (total n=4). VC, virus control. CC, cell control. (c) Virus yield reduction assay with Rupintrivir and Vapendavir. Viral RNA in supernatants was quantified at 72 hpi. Data are mean ± SD (n=3). (d, e) Proof-of-concept inter-operator assay reproducibility across >300 plates containing pre-spotted reference compounds in assays with RV-A16, RV-B14, and RV-C15a-mGL. (f) Representative images of RV-C11a-mGL and RV-C41a-mGL infection and antiviral assays with Rupintrivir (n=3).

Assay performance was evaluated with three reference compounds representing distinct antiviral mechanisms. Rupintrivir is a pan-entero/-rhino 3C protease inhibitor and its activity against RV-C15 has been confirmed in HeLa cells using a replicon system (8) and in human airway epithelial (HAE) culture (8, 25). BF738735, a PI4KB inhibitor, has not been directly tested against RV-C, but the related PI4KB inhibitor PIK93 was reported to inhibit multiple RV-C replicons and RV-C infection in HAE culture (8). Pirodavir, a broad-spectrum capsid binder against RV-A and RV-B, is ineffective against RV-C due to its narrower hydrophobic capsid pocket (8, 25). In our assay, Rupintrivir and BF738735 inhibited RV-C15a-mGL replication with IC_50_ values of 9 and 14 nM, respectively (Fig. 2b), comparable with their reported activity against rhinoviruses. As expected, Pirodavir had no effect.

To further validate the system, virus yield reduction assays were performed with Rupintrivir and Vapendavir, another capsid binder. Viral RNA in cell culture supernatants was quantified at 72 hpi (Fig. 2c). Rupintrivir treatment reduced viral RNA by ∼3-log_10_ compared with untreated cells, whereas Vapendavir showed no effect. Together, these results confirm that the assay reliably detects potent inhibitors while excluding compounds with RV-A/B–specific activity, thereby demonstrating biological specificity.

Next, a proof-of-concept study was conducted to assess assay robustness using pre-spotted plates containing the three reference compounds (BF738735, Pirodavir and Rupintrivir), tested across 360 plates by two independent operators (Fig. 2d, S4c, and S4d). In parallel, the prototypic strains of RV-A16 (species A) and RV-B14 (species B) were evaluated in the same cell line, enabling a unified framework covering all three RV species. The RV-C15 assay achieved a mean Z′-factor of 0.80 ± 0.10, exceeding the 0.5 threshold for HTS suitability, demonstrating an excellent assay window and reproducibility. Comparable Z’ values were obtained for RV-A16 (0.88 ± 0.06) and RV-B14 (0.89 ± 0.05) under identical conditions. In addition, low inter-operator coefficients of variation (CV) (Fig. S4c) and high concordance in IC_50_ values for across three RV species (Fig. 2e and S4d) further confirmed assay reproducibility. Together, these data demonstrate the flexibility of the platform and its applicability beyond RV-C.

Finally, to assess broader applicability, the assay was extended to two additional RV-C types (RV-C11 and RV-C41 reporter viruses). Although their replication efficiency was slightly lower than RV-C15, both produced sufficient fluorescence signals for HCI. Rupintrivir inhibition profiles were similar to those observed for RV-C15, demonstrating that the platform can be directly applied across multiple RV-C types (Fig. 2f).

### Miniaturization to 384-Well Format and HTS Validation

To enable high-throughput compound screening, the optimized 96-well antiviral assay was miniaturized to a 384-well format. Seeding densities from 1 000 to 4 000 cells per well were tested in combination with a three-point serial dilution of RV-C15a-mGL to determine the most robust conditions (Fig. S5a). Across all tested combinations, including virus dilution from 1/30 to 1/2430, the assay consistently achieved a Z′-factor above 0.7. Two conditions (highlighted by white border, Fig. S5a) additionally supported infection rates above 50% and cell viability above 70%, parameters considered optimal for HTS applications. Based on these results, an MOI of 0.0005 (equivalent to 1/2500 dilution) with 3 000 cells per well was selected for subsequent antiviral assays.

Assay robustness in the 384-well format was further validated across the three RV species using pre-spotted plates containing Rupintrivir, BF738735, and Pirodavir (Fig. 3a). Across more than 140 independent plates, mean Z′-factor of 0.71 ± 0.06 (n=141), 0.73 ± 0.08 (n=141), and 0.79 ± 0.07 (n=150) were obtained for RV-A, RV-B, and RV-C, respectively, comparable to values obtained in the 96-well format (Fig. 3b). IC_50_ values were highly consistent across plates (Fig. 3c and S5b) and comparable to those from the 96-well assay (Fig. 3d), confirming that miniaturization did not compromise assay reproducibility.

**Figure 3.**
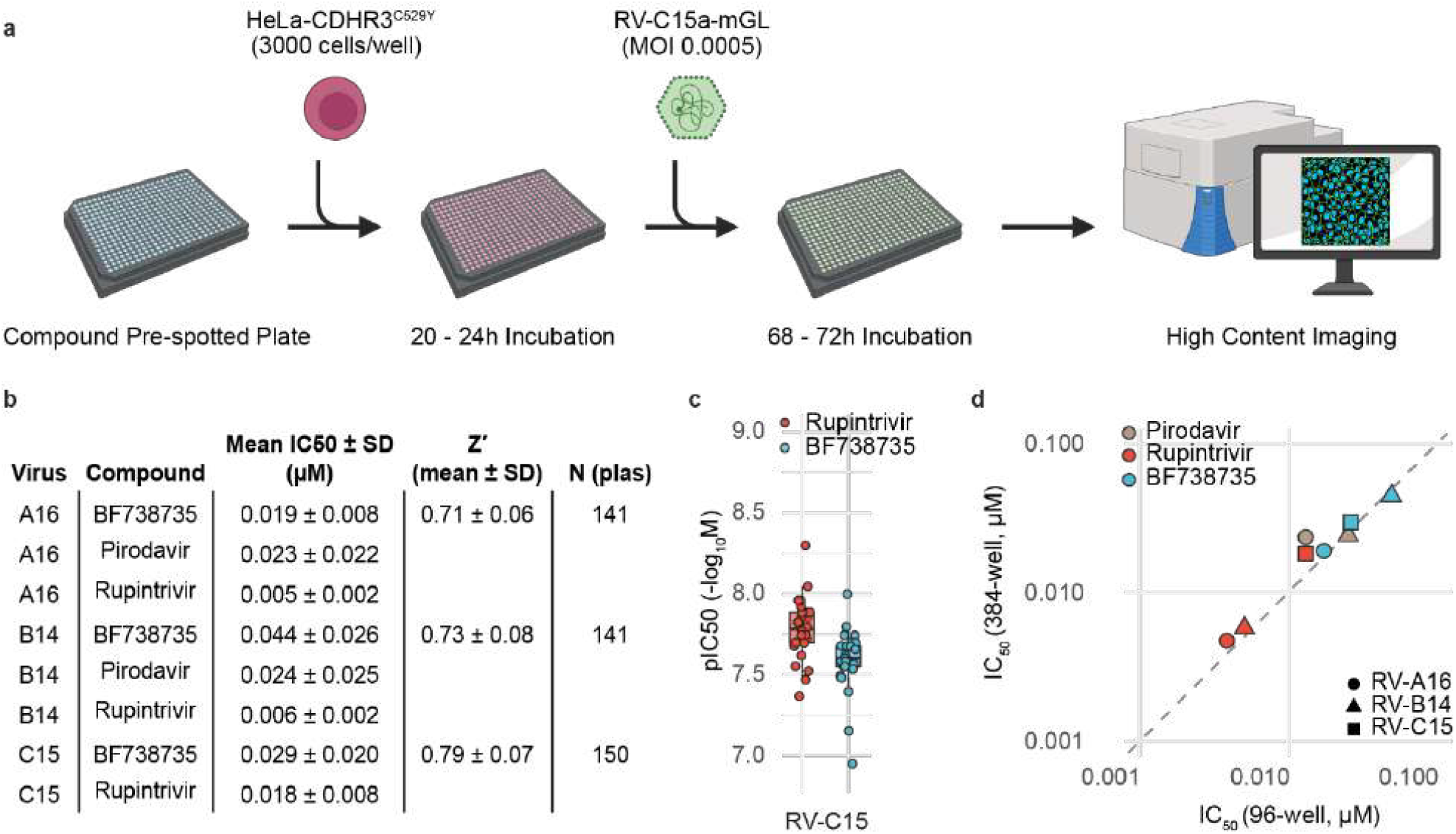
Miniaturization of the RV-C15a-mGL assay to 384-well format. (a) Schematic of 384-well format antiviral assay procedure. Created with BioRender. (b) Assay robustness in 384-well format across RV-A16, RV-B14, and RV-C15a-mGL assays using >100 pre-spotted plates with reference compounds. (c) Distribution of pIC_50_ values (-log_10_ IC_50_, M) for BF738735 and Rupintrivir in the RV-C15a-mGL assay. Each dot represents an individual run; boxes indicate median, interquartile range (IQR), and whiskers (1.5x IQR).

### Preliminary High-Throughput Screening and Hit Validation

To evaluate the applicability of the optimized 384-well RV-C15a-mGL assay for high-throughput screening, we performed a proof-of-concept screen of a medium-sized library comprising 10 240 compounds, curated for HTS suitability by removing pan-assay interference compounds (PAINS) and ensuring structural diversity and chemical integrity of compounds (Fig. 4a). Compounds were tested at a single concentration of 10 µM, with virus-only and uninfected cell controls included for normalization. Reference compounds (Rupintrivir, BF738735, and Pirodavir) were tested in parallel. As expected, Rupintrivir and BF738735 completely inhibited virus replication, although BF738735 also slightly reduced cell viability at this concentration, whereas Pirodavir showed no measurable effect on RV-C15 replication (Fig. 4b). Assay performance remained robust, with Z’-factors consistently above 0.8 across 32 plates (Fig. 4c). From the primary screening, 19 compounds reduced mGL fluorescence by ≥60% while maintaining ≥90% cell viability. These compounds were re-tested in mini dose–response assays (0.2–25 µM, four replicates), from which IC_50_, CC_50_, and selectivity index (SI = CC_50_/IC_50_) values were determined (Table S1, Fig. 4d, 4e, and S6). Compounds with SI > 7 were classified as confirmed hits, yielding five validated RV-C15 antiviral candidates.

**Figure 4.**
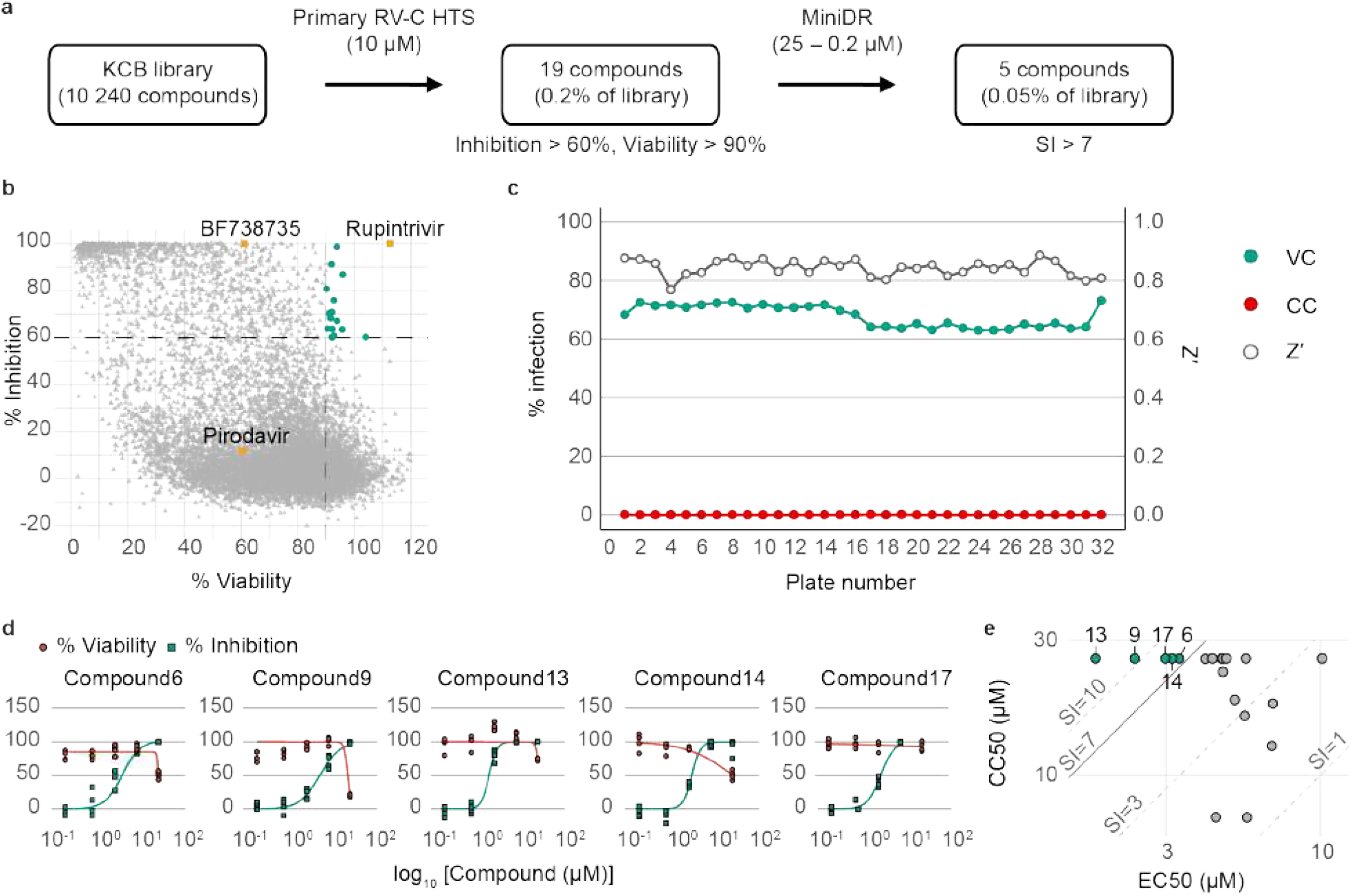
Preliminary high-throughput screening of a 10 240-compound library. (a) Workflow of RV-C15a-mGL HTS campaign in 384-well format. (b) Scatterplot summarizing primary screening at 10 µM and three reference compounds BF738735, Pirodavir, and Rupintrivir (yellow). Compounds with inhibition > 60% and viability > 90% are highlighted in green. (c) Assay performance across 32 HTS plates. VC and CC are plotted against the left Y-axis in % infection, while Z’ is plotted against the right Y-axis. VC, virus control. CC, cell control. (d) Inhibition and viability curves for five top hit compounds from mini dose-response assays (0.2 – 25 µM, n=4). (e) Distribution of IC_50_, CC_50_, and selective index (SI = CC_50_/IC_50_) values for 19 confirmed hits from mini dose-response assays. Five top hits (SI > 7) are highlighted in green.

Although detailed characterization of these compounds lies beyond the scope of this study, these proof-of-concept results demonstrate the scalability of the assay and establish its utility as a high-throughput screening platform for antiviral discovery.

## Discussion

RV-C is an important cause of asthma exacerbations and severe lower respiratory infections, yet its study has been hampered by the absence of robust *in vitro* models. The identification of CDHR3 as the RV-C receptor enabled virus replication in receptor-engineered HeLa cells (14), but this model was not adapted for systematic drug discovery. Here, we established and validated a high-throughput screening (HTS) platform for RV-C, integrating receptor-engineered cells, phenotypic clone selection, and a brighter and genetically stable mGreenLantern reporter virus.

A key strength of this work is the phenotypic selection strategy used to identify the most permissive and stable monoclonal clones, rather than relying solely on receptor overexpression alone. This approach revealed that receptor abundance is not the sole determinant of RV-C susceptibility. In contrast to the clear link between ACE2 expression and SARS-CoV-2 entry (26), clones with the highest CDHR3 levels were not necessarily the most susceptible to RV-C infection (Fig. 1, S2, and S3). Moreover, several initially permissive clones lost susceptibility over time despite sustained high receptor expression, suggesting that permissiveness depends on a combination of receptor availability and other cellular factors. Together, these findings emphasize that productive RV-C replication is orchestrated by multiple determinants beyond receptor density, highlighting the importance of functional phenotypic selection and long-term monitoring of engineered systems. More research is required to identify which and how these underlying factors contribute to supporting RV-C infection.

Through this process, mono29 was established as a robust and stable cell line supporting consistent infection across passages, providing a reliable foundation for assay development. The optimized assays in both 96-(Fig. 2 and S4) and 384-well (Fig.3 and S5) formats consistently yielded Z′ factors above 0.7 and reproducible IC_50_ values across independent operators, confirming the assay’s robustness and scalability. Importantly, the same framework supported RV-A16 and RV-B14, and was directly adaptable to additional RV-C types (C11 and C41), highlighting its versatility across rhinovirus species and providing a unified system for side-by-side evaluation of virus replication and inhibitor efficacy.

A successful pilot screen of 10 240 compounds further demonstrated the scalability and reliability of the platform in a true high-throughput format, yielding several confirmed RV-C15 inhibitors with selective antiviral activity. While detailed characterization of these candidates lies beyond the scope of this study, their identification highlights the readiness of this assay for integration into drug discovery pipelines. This system therefore bridges the gap between physiologically relevant but low-throughput airway cultures (10, 25) and high-throughput yet less representative models such as replicons (8), offering a biologically relevant and scalable platform for systematic antiviral evaluation.

Beyond its application to antiviral testing, this platform also opens new opportunities to investigate fundamental aspects of RV-C biology that have so far remained underexplored. The platform allows controlled analyses of replication kinetics and infection dynamics, and enables detailed studies of CDHR3-mediated virus entry mechanism. As the canonical function of CDHR3 remains poorly understood, such studies may also provide insights into the physiological role of this receptor and its interaction with the virus. In addition, the system can be used to investigate previously identified proviral host factors, such as STING and lipid-associated pathways (27) in a biologically relevant setting that supports the full replication cycle. Likewise, antiviral host-factors or asthma-associated factors beyond CDHR3 (15) can be examined to better understand their contribution to RV-C susceptibility and disease severity.

In conclusion, we established a validated, scalable, and reproducible HTS-compatible platform for RV-C research. This platform not only enables systematic identification of RV-C antivirals but also provides a unified experimental framework applicable across rhinovirus species. Beyond drug discovery, it offers a foundation for investigating the full replication cycle of RV-C, virus-host interactions, and the effects of host-targeted inhibitors. Together, these advances address a longstanding limitation in RV-C research and open new opportunities for both therapeutic development and mechanistic studies of rhinovirus biology.

## Materials and Methods

### Detailed materials and methods are provided in the SI Appendix

#### Cells

HeLa-Rh (28) and HEK293T (ATCC, CRL-3216), cells were maintained in Dulbecco’s Modified Eagle Medium (DMEM) (Gibco) supplemented with 10% fetal bovine serum (FBS) (HyClone) and 1% penicillin-streptomycin (10,000 U/mL, Gibco). HeLa cells stably expressing CDHR3^C529Y^ were generated by lentiviral transduction and selected in DMEM with 10% FBS, 1% pen-strep, and 2.5 µg/mL blasticidin. All cells were maintained at 37 °C with 5% CO_2_, and were routinely tested negative for mycoplasma.

#### Viruses

RV-A16 and RV-B14 were derived from infectious clones pR16.11 (ATCC VRMC-8) and pWR3.26 (ATCC VRMC-7) and produced in HeLa-Rh cells. A fluorescent reporter virus (pRV-C15a-mGL; strain W10) was engineered by adding adaptive mutations (VP1-T125K, 3A-E41K) by multisite mutagenesis and by inserting syntheized mGreenLantern (GenBank OP373690.1) between VP1 and 2A. Reporter clones for RV-C11 (strain CL-170085, EU840952.2) and RV-C41 (isolate 2536, KF958311.1) were generated using pUC19-CMV/T7 backbone, and 3A-E41K mutation was added. All RV-Cs were produced in polyclonal HeLa-CDHR3^C529Y^ cells. Virus titer was determined by end-point dilution assay using high-content imaging (HCI) with Thermo Fisher CX5 platform.

#### Monoclonal clone selection

Single-cell sorting was performed using Pala Cell Sorter and Single Cell Dispenser (bio-techne) into 96-well plates. Expanded monoclonal cells were screened for RV-C15a-mGL permissiveness using a fixed virus dose in triplicate. Cells were fixed at 3 dpi with 4% paraformaldehyde (PFA), stained with DAPI, and quantified by HCI. Clones with >50% infection in two out of three replicates were classified as permissive and validated by flow cytometry.

#### Growth kinetics

HeLa-CDHR3^C529Y^ mono29 cells were seeded in 96-well plates in infection medium (DMEM with 2% FBS and 30 mM MgCl_2_), and were grown for 20 – 24h at 37 °C. Cells were inoculated with RV-C15a-mGL at different MOIs and incubated at 32 °C for 24, 48, or 72 h. Cells were fixed with 4% PFA, stained with DAPI, and quantified by HCI. Z′ factors were calculated for each condition.

#### Compounds

Rupintrivir (Sigma-Aldrich PZ0315), BF738735 (MedChemExpress HY-U00426), Pirodavir (Sigma-Aldrich SML1818), and Vapendavir (Cayman 30546) were prepared as 10 mM DMSO stocks. Compounds were dispensed in assay-ready plates using I.DOT or Echo. The screening library comprised 10 240 small molecules provided by the Korea Chemical Bank (KCB), representing a diversity set curated to be HTS-friendly by removing PAINS and including compounds with high structural diversity, favorable drug-like properties, and verified purity.

#### Pilot antiviral assay

HeLa-CDHR3^C529Y^ mono29 were seeded at 7,500 cells/well in 96-well plates. Compound dilution series was added to the cells. Cells were infected with RV-C15a-mGL (MOI 0.002). At 3 dpi, cells were fixed, stained with DAPI, and quantified by HCI. IC_50_ and CC_50_ values were determined by nonlinear regression in GraphPad Prism v10.6.1.

#### Virus yield reduction assays

HeLa-CDHR3^C529Y^ mono29 were infected with RV-C15a-mGL (MOI 0.01) for 1h, washed twice with PBS, and incubated with compound-containing medium. Supernatants were harvested at 72 hpi, and viral RNA was extracted (MagNA Pure 96, Roche) and quantified by RT-qPCR (TaqMan Fast Virus 1-Step, Thermo Fisher) with corresponding primers and probe (Table S2).

#### 96-well assay validation with reference compounds

Assays were performed with pre-spotted plates (96-well) under conditions described above. IC_50_, CC_50_, Z′, and inter-operator CVs were determined using Dotmatics and R v4.4.2.

#### Assay miniaturization to 384-well format and automation

Assays were scaled to 384-well plates with automated dispensing (MultiDrop Combi, Thermo Fisher; MultiFlo Fx with BioStack, Agilent). Plates were fixed at 3 dpi, washed, stained with DAPI, and analyzed by HCI.

#### High-throughput screening

The KCB library was screened in 384-well format at 10 µM. Primary hits were defined as ≥60% reduction of GFP signal with ≥90% viability. Hits were retested in mini dose–response assays (0.2–25 µM). Screening data were processed in Genedata Screener®. Dose–response curves were re-generated in GraphPad Prism for visualization, and selective index (SI) values were calculated in R.

#### High-content imaging and analysis

HCI was performed using CellInsight CX5 with Orbitor RS2 robotic platform and in-house adapted SpotDetector v4.1 protocol. Images were acquired in DAPI (Ex/Em 386/440 nm) and FITC (485/521 nm) channels. For RV-A16/B14, four fields per well were imaged with a 4x objective; for RV-Cs, nine fields per well with a 10x objective. Data were normalized to virus control (VC) and cell control (CC).

#### Statistical analysis

Data were analyzed in GraphPad Prism v10.6.1, R v4.4.2, Dotmatics, and Genedata Screener v20.0.7. Z′ and CVs were calculated as described (29) in R. Spearman correlation was used to assess relationships between infection efficiency and CDHR3 or HS expression.

## Acknowledgment

HeLa Rh was kindly provided by K. Andries (Janssen Pharmaceutica, Belgium). We thank Jasper Rymenants for technical assistance with the experiments involving human epithelial cell cultures, and Dirk Jochmans for supervision of these experiments. We also thank Nelleke Cloet for preparing and providing the pre-spotted assay plates, and Hugo Klaassen for supervising this work.

The high-throughput screening campaign was supported by the National Research Foundation of Korea (NRF) grant funded by the Korean Ministry of Science and ICT (No. RS-2024-00432287).

## Author Contributions

H.L., Y.A.A., K.D., P.L., S.H., J.N. and H.J.T designed research; Y.A.A., C.S., T.R., J.P., M.R., and N.T. performed research; C-S.Y., N-C.C., and S.B.H. contributed library and compounds; H.L., Y.A.A., C.S., T.R., and P.L. analyzed data; and H.L., Y.A.A., J.N. and H.J.T. wrote the paper.

## References

1. S. Lau, et al., Clinical Features and Complete Genome Characterization of a Distinct Human Rhinovirus (HRV) Genetic Cluster, Probably Representing a Previously Undetected HRV Species, HRV-C, Associated with Acute Respiratory Illness in Children. J. Clin. Microbiol. 45, 3655–3664 (2007).

2. N. Renwick, et al., A Recently Identified Rhinovirus Genotype Is Associated with Severe Respiratory-Tract Infection in Children in Germany. J. Infect. Dis. 196, 1754–1760 (2007).

3. Wai-ming Lee, et al., A Diverse Group of Previously Unrecognized Human Rhinoviruses Are Common Causes of Respiratory Illnesses in Infants. PLoS ONE 2 (2007).

4. P. McErlean, et al., Distinguishing Molecular Features and Clinical Characteristics of a Putative New Rhinovirus Species, Human Rhinovirus C (HRV C). PLoS ONE 3 (2008).

5. E. Miller, et al., Human rhinovirus C associated with wheezing in hospitalised children in the Middle East. J. Clin. Virol. Off. Publ. Pan Am. Soc. Clin. Virol. 46 1, 85–9 (2009).

6. N. Khetsuriani, et al., Novel Human Rhinoviruses and Exacerbation of Asthma in Children. Emerg. Infect. Dis. 14, 1793–1796 (2008).

7. Y. A. Bochkov, et al., Molecular modeling, organ culture and reverse genetics for a newly identified human rhinovirus C. Nat. Med. 17, 627–632 (2011).

8. C. Mello, et al., Multiple Classes of Antiviral Agents Exhibit In Vitro Activity against Human Rhinovirus Type C. Antimicrob. Agents Chemother. 58, 1546–1555 (2014).

9. W. Hao, et al., Infection and Propagation of Human Rhinovirus C in Human Airway Epithelial Cells. J. Virol. 86, 13524–13532 (2012).

10. C. Tapparel, et al., Growth and characterization of different human rhinovirus C types in three-dimensional human airway epithelia reconstituted in vitro. Virology 446, 1–8 (2013).

11. T. B. Gagliardi, et al., Rhinovirus C replication is associated with the endoplasmic reticulum and triggers cytopathic effects in an in vitro model of human airway epithelium. PLoS Pathog. 18 (2021).

12. C. Li, et al., Human respiratory organoids sustained reproducible propagation of human rhinovirus C and elucidation of virus-host interaction. Nat. Commun. 15, 10772 (2024).

13. M. Essaidi-Laziosi, et al., Rhinovirus C replication and host interactions in differentiated airway epithelia. Viruses 15, 317 (2023).

14. Y. A. Bochkov, et al., Cadherin-related family member 3, a childhood asthma susceptibility gene, mediates rhinovirus C binding and replication. Proc Natl Acad Sci U A 112, 5485–5490 (2015).

15. K. Bønnelykke, et al., A genome-wide association study identifies CDHR3 as a susceptibility locus for early childhood asthma with severe exacerbations. Nat. Genet. 46, 51–55 (2014).

16. K. Bønnelykke, et al., Cadherin-related family member 3 genetics and risk of severe asthma exacerbations by rhinovirus C. J Allergy Clin Immunol 139, 587–596 (2017).

17. T. F. Griggs, et al., Rhinovirus C replication in HeLa cells expressing cadherin-related family member 3. J Virol 91, e00045–17 (2017).

18. J. L. Everman, et al., Functional genomics of CDHR3 confirms its role in HRV-C infection and childhood asthma exacerbations. J. Allergy Clin. Immunol. 144, 962–971 (2019).

19. S. Basnet, et al., CDHR3 Asthma-Risk Genotype Affects Susceptibility of Airway Epithelium to Rhinovirus C Infections. Am. J. Respir. Cell Mol. Biol. 61, 450–458 (2019).

20. D. Olszewski, et al., High-content, arrayed compound screens with rhinovirus, influenza A virus and herpes simplex virus infections. Sci. Data 9 (2022).

21. W. Chiu, et al., Development of a robust and convenient dual-reporter high-throughput screening assay for SARS-CoV-2 antiviral drug discovery. Antiviral Res. 210, 105506 (2023).

22. Y. A. Bochkov, et al., Mutations in VP1 and 3A proteins improve binding and replication of rhinovirus C15 in HeLa-E8 cells. Virology 499, 350–360 (2016).

23. B. C. Campbell, et al., mGreenLantern: a bright monomeric fluorescent protein with rapid expression and cell filling properties for neuronal imaging. Proc. Natl. Acad. Sci. 117, 30710–30721 (2020).

24. A. K. Patick, et al., In Vitro Antiviral Activity of AG7088, a Potent Inhibitor of Human Rhinovirus 3C Protease. Antimicrob. Agents Chemother. 43, 2444–2450 (1999).

25. B. Boda, et al., Antiviral drug screening by assessing epithelial functions and innate immune responses in human 3D airway epithelium model. Antiviral Res. 156, 72–79 (2018).

26. S. Kazemi, et al., Variations in Cell Surface ACE2 Levels Alter Direct Binding of SARS-CoV-2 Spike Protein and Viral Infectivity: Implications for Measuring Spike Protein Interactions with Animal ACE2 Orthologs. J. Virol. 96, e00256–22 (2022).

27. K. McKnight, et al., Stimulator of interferon genes (STING) is an essential proviral host factor for human rhinovirus species A and C. Proc. Natl. Acad. Sci. U. S. A. 117, 27598–27607 (2020).

28. D. Roymans, et al., Therapeutic efficacy of a respiratory syncytial virus fusion inhibitor. Nat. Commun. 8, 167 (2017).

29. J. H. Zhang, T. D. Chung, K. R. Oldenburg, A Simple Statistical Parameter for Use in Evaluation and Validation of High Throughput Screening Assays. J. Biomol. Screen. 4, 67–73 (1999).

