## Supporting Information for "A high-throughput cell-based platform for Rhinovirus C research and antiviral drug discovery"

**This PDF file includes:**

Materials and Methods
Figures S1 to S6
Tables S1 to S3
SI References

**Supporting Information**

**Materials and Methods**

**Plasmid construction and mutagenesis.**

All primers and synthetic DNA are listed in Table S2 and S3.

*pLenti-CDHR3$^{C529Y}$-IRES-Blast*

The pLenti-CDHR3$^{C529Y}$-IRES-Blast construct was generated by replacing Cas9 in pLenti-Cas9-2A-Blast (Addgene #73310) with PCR-amplified human CDHR3 cDNA encoding the Y529 variant (from pGL4-CMV-FLAG-HA-CDHR3C529Y, kindly provided by Maarten Jacquemyn).

*pRV-C15a, pRV-C15a-mGL*

Full-length RV-C15 (strain W10, GenBank GU219984.1) was synthesized from three gBlocks and one Ultramer (IDT) including a T7 promoter and flanking pUC19 overlaps, and assembled using NEBuilder HiFi. Adaptive mutations (VP1-T125K, 3A-E41K) were introduced by multisite mutagenesis, yielding clone pRV-C15a. A fluorescent reporter virus (pRV-C15a-mGL) was engineered by inserting synthesized mGreenLantern (GenBank OP373690.1) between VP1 and 2A, as previously described for RV-C15-GFP (1).

*pRV-C11a-mGL and pRV-C41a-mGL*

Reporter clones for RV-C11 (strain CL-170085, EU840952.2) and RV-C41 (isolate 2536, KF958311.1) were generated in a similar manner but using pUC19-CMV/T7 as the backbone.

*pUC19-CMV/T7*

Based on a universal cloning system previously reported (2), we generated pUC19-CMV/T7 to streamline future RV-C infectious clone generation. The CMV promoter and T7 promoter were amplified from the pcDNA3.1 vector. Hammerhead ribozyme, hepatitis delta ribozyme, and SV40 poly(A) sequences were synthesized. All fragments were assembled using NEBuilder HiFi.

All constructs were confirmed by whole plasmid sequencing.

**Virus production.** RV-A16 (VRMC-8) and RV-B14 (VRMC-7) infectious clones were obtained from ATCC and linearized with SacI and MluI, respectively. All RV-C infectious clones were linearized with XmaI. The linearized DNA was transcribed *in vitro* using T7 RiboMAX system (Promega), and the resulting RNA was purified with the NucleoSpin RNA Mini kit (Macherey-Nagel). RNA was transfected into HeLa-Rh cells (RV-A16 and RV-B14) or polyclonal HeLa-CDHR3$^{C529Y}$ cells (RV-Cs) using Lipofectamine™ MessengerMAX™ (ThermoFisher), and cells were incubated at 32°C for 3 days. After three freeze-thaw cycles, cell lysates were clarified by low-speed centrifugation and filtered through a 0.2 µm membrane. Virus titers were determined by end-point titration and expressed as 50% tissue culture infectious dose ($TCID_{50}$).

**Generation of HeLa cells stably expressing CDHR3$^{C529Y}$.** HEK293T cells were co-transfected with pLenti-CDHR3$^{C529Y}$-IRES-Blast, psPAX2, and pMD2.G (1:1:0.5 ratio) using FuGENE® 4K. Supernatants were harvested 48 hours post-transfection, clarified, and used to transduce HeLa-Rh cells in the presence of 8 μg/mL polybrene. Stable cells were selected with 2.5 μg/ml blasticidin until all non-transduced controls had died (10 – 14 d).

**Immunostaining.** Polyclonal HeLa-CDHR3$^{C529Y}$ cells were grown in 96-well plates. Subconfluent cells were infected with RV-C15a-mGL for 1 h at 32°C. Inoculum was removed and fresh infection medium was added. At 48 hpi, cells were fixed with 4% PFA for 15 min at room temperature. After permeabilization with 0.1% Triton X-100 in PBS for 5 min, cells were incubated with anti-dsRNA antibody (Jena Bioscience, RNT-SCI-10010200; 1:1000) in 1% bovine serum albumin (BSA) in PBS for 2 h at room temperature. After washing twice with PBS, cells were incubated with goat anti-mouse Alexa647 (Invitrogen™ A21235; 1:500) for 1 h at room temperature. Nuclei were stained with DAPI. Images were acquired by HCI.

**RT-PCR for RV-C15a-mGL stability.** Viral RNA was isolated from passages P0 to P2 of RV-C15a-mGL produced from polyclonal HeLa-CDHR3$^{C529Y}$ cells using NucleoSpin RNA Virus kit (Macherey-Nagel). RNA was used as template for RT-PCR with qScript™ XLT One-Step RT-PCR Kit (Quantabio) under the following conditions: 20 min at 55°C, 3 min at 94°C, 40 cycles of 30 sec at 94°C, 45 s at 60°C, 72 sec at 72°C, and final extension of 10 min at 72°C). One-fifth of each PCR product was analyzed by gel electrophoresis, and sequences were verified by Sanger sequencing.

**RV-C15a growth curve and antiviral assay in human primary epithelial air-liquid interface (ALI) culture.** Human bronchial epithelial (HBE; MucilAir EP01) and nasal airway epithelial (HNE; MucilAir EP02) cells (Epithelix) were obtained in an ALI format. Upon arrival, inserts were washed with pre-warmed PBS and maintained in MucilAir medium (Epithelix EP04MM) at 37 °C and 5% CO2 for at least four days before use. For infection, apical surfaces were washed twice with pre-warmed medium to remove accumulated mucus and placed in new 24-well plates containing fresh medium. Cultures were infected apically with 100 μl of RV-C15a for 1 h at 35°C, washed, and incubated at 35°C. For antiviral assays, cultures were pretreated basolaterally with compound-containing medium at different concentrations for 1 h before apical infection with 3.6 TCID50 per insert. After 1 h, inoculum was removed, and compound-containing medium in the basolateral side was refreshed on day 2 after infection. Apical surfaces were washed at designated time points, and washes were subjected to Cells-to-cDNA II kit (Invitrogen™ AM1722). RV-C15 copy number was determined by RT-qPCR using iTaq Universal Probes One-Step Kit (BIO-RAD).

**Flow cytometry.** A total of $10^5$ cells per well were seeded into a U-bottom 96-well plate. Cells were detached using Versene (Thermo Fisher Scientific) rather than trypsin to prevent shedding of surface proteins. Plates were centrifuged for 5 min at 500 x g in a pre-cooled centrifuged at 4 °C, and supernatants were carefully discarded. After each centrifugation, the plate was briefly slid over a vortex platform to resuspend the cell pellet in the residual fluid. Cells were washed with 200 μL phosphate-buffered saline

(PBS) supplemented with 2.5 mM EDTA (PBS-EDTA), followed by centrifugation and supernatant removal as described above. Cells were then incubated for 20 min at room temperature (RT) with 50 µL of a primary antibody mix containing rabbit polyclonal anti-CDHR3 (1:50; Atlas Antibodies, HPA011218) and mouse monoclonal anti-heparan sulfate (IgM, 1:400; AMSBIO, 370255-S) prepared in FACS buffer (PBS supplemented with 2% FBS and 2.5 mM EDTA). After washing, cells were incubated for 20 min at RT in the dark with 50 µL of a secondary antibody mix containing Brilliant Violet 421™ donkey anti-rabbit IgG (BioLegend, Cat. No. 406410) and Brilliant Violet 711™ anti-mouse IgM (BioLegend, Cat. No. 406516). Cells were washed again, resuspended in 100 µL PBS-EDTA, and fixed by addition of 100 µL PBS containing 8% PFA (final concentration 4%) for 15 min at RT. Following fixation, cells were washed once more and resuspended in 200 µL FACS buffer. Samples were transferred to flow cytometry tubes and analyzed on a LSRFortessa™ X-20 flow cytometer (BD Biosciences). Flow cytometry data were analyzed using FlowJo software (v10.9; BD Biosciences).

**384-well format assay condition optimization.** HeLa-CDHR3$^{C529Y}$ mono29 cells were seeded at densities ranging from 1,000 to 4,000 cells/well in 384-well plates. Cells were infected with a 10-point, 3-fold serial dilution of RV-C15a-mGL. Optimal assay conditions were defined as those yielding robustness assay performance (Z' > 0.7, % infection > 50, % viability > 70) at 3 dpi.

### Figures

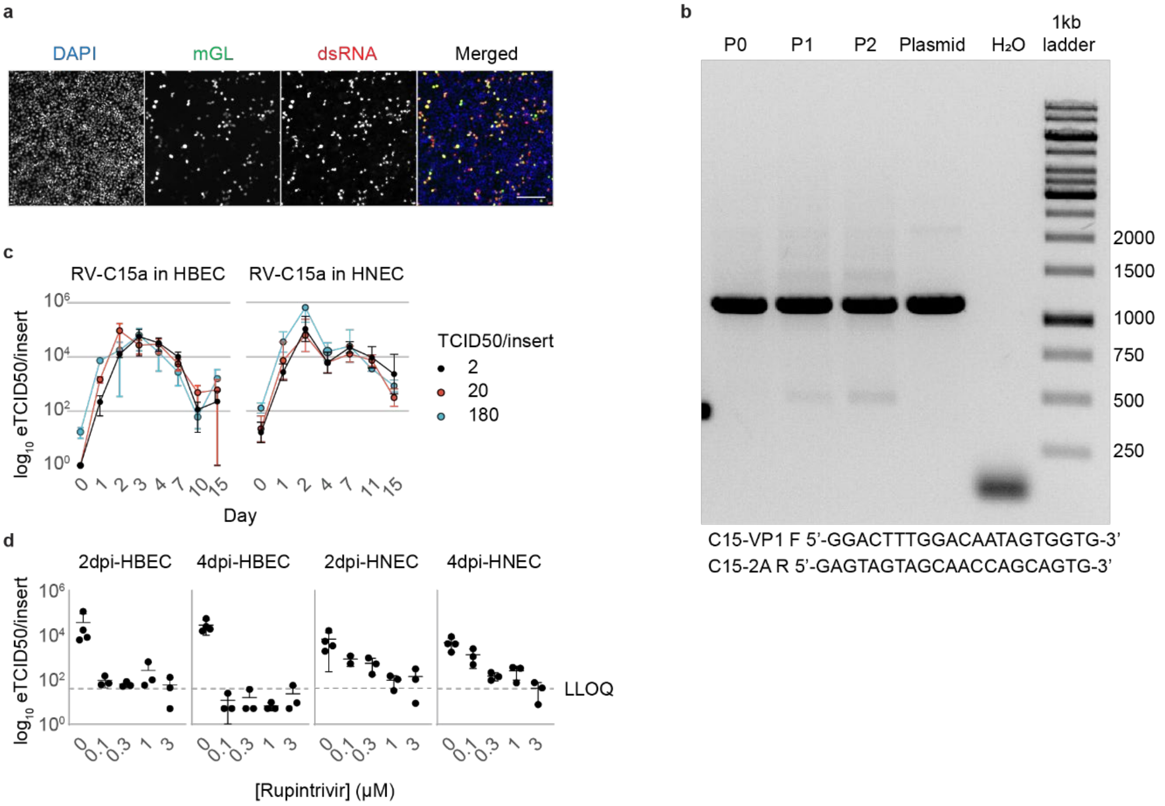


**Figure S1. Characterization of RV-C15a and RV-C15a-mGL reporter virus.** (a) Representative images showing mGL signal and dsRNA staining in RV-C15a-mGL infected polyclonal HeLa-CDHR3$^{C529Y}$ at 48 hours post infection (hpi). Scale bar is 200 μm. (b) RT-PCR fingerprinting of viral RNA confirming mGL insert stability across two passages. Viral RNA was isolated from passages P0 to P2 of RV-C15a-mGL produced from polyclonal HeLa-CDHR3$^{C529Y}$ cells. (c) Replication of RV-C15a in human bronchial epithelial cells (HBEC) and human nasal airway epithelial cells (HNEC) at air-liquid interface assessed by RT-qPCR. Values represent mean values of triplicates ± standard deviation. (d) Inhibition of RV-C15a by Rupintrivir in HBEC and HNEC cultures. Values represent mean values of triplicates ± standard deviation, except for untreated (n=4). LLOQ, lower limit of quantification.

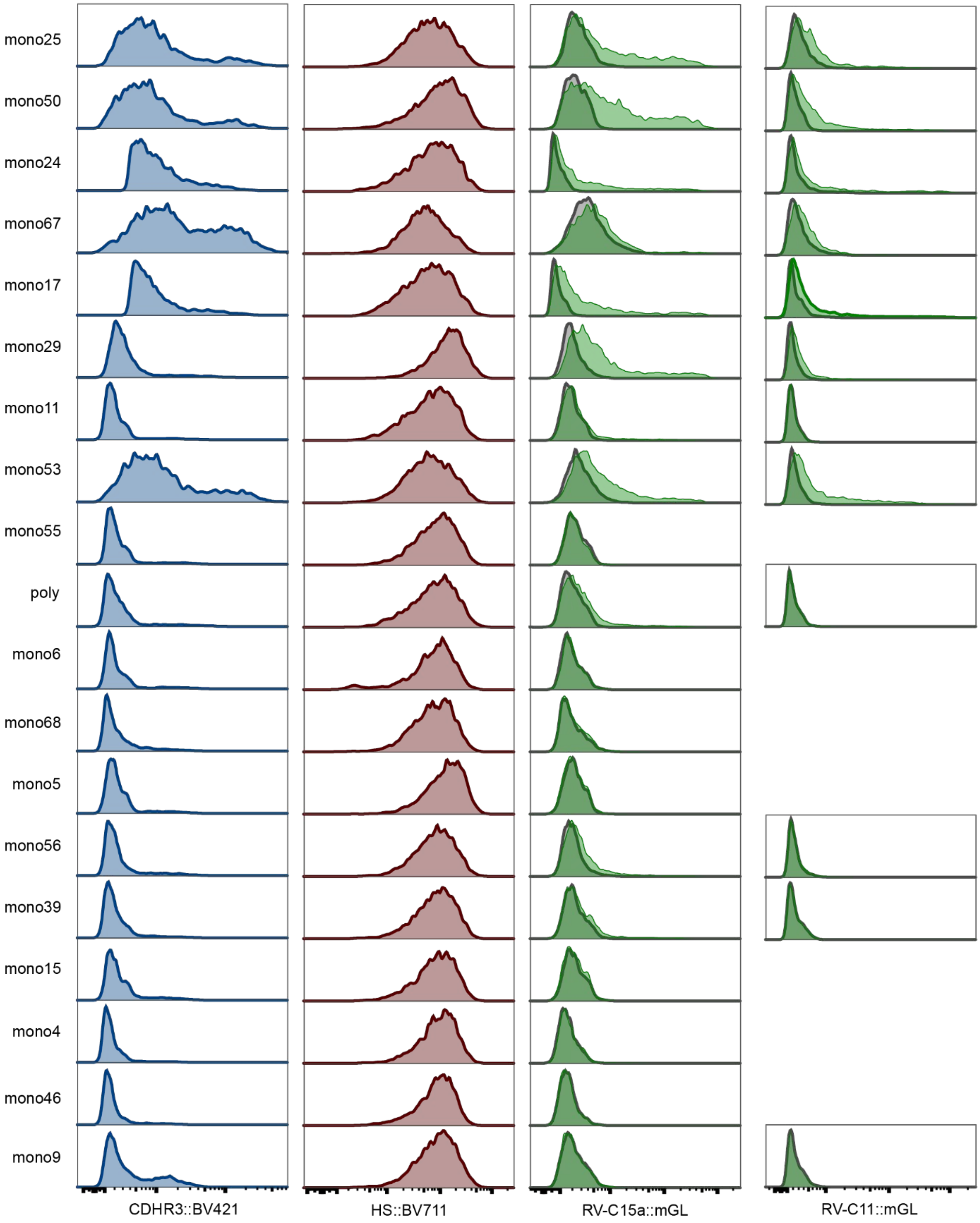


**Figure S2. Flow cytometry characterization of monoclonal cell clones.** Representative histograms showing CDHR3 (blue) and heparan sulfate (HS, red) surface expression. Virus infection efficiency was measured by mGreenLantern fluorescence intensity.

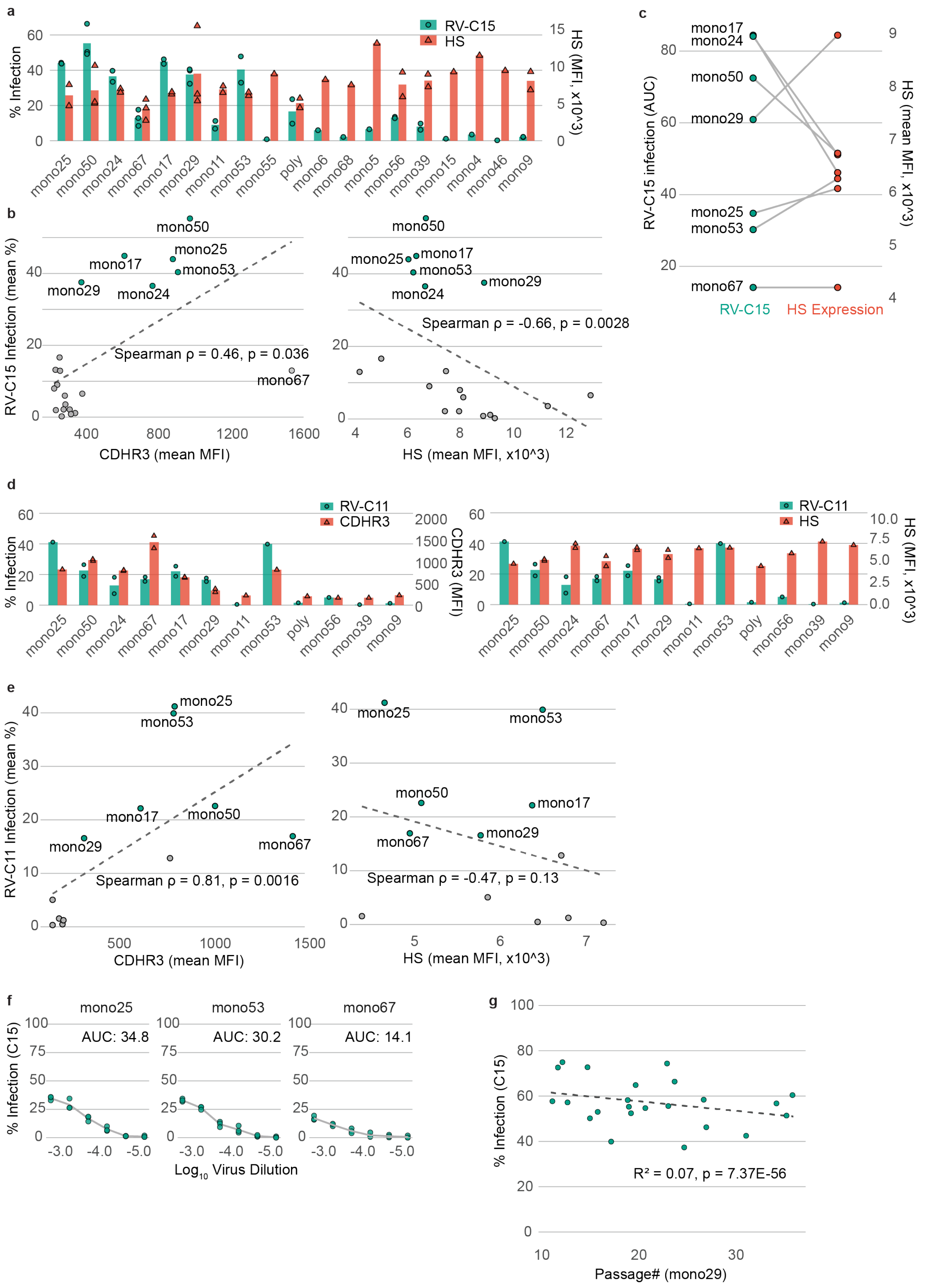
a
% Infection
HS (MFI, x10^3)
RV-C15
HS
mono25 mono50 mono24 mono67 mono17 mono29 mono11 mono53 mono55 poly mono6 mono68 mono5 mono56 mono39 mono15 mono4 mono46 mono9
b
RV-C15 Infection (mean %)
mono50
mono25
mono17
mono53
mono29
mono24
Spearman ρ = 0.46, p = 0.036
mono67
CDHR3 (mean MFI)
mono25
Spearman ρ = -0.66, p = 0.0028
HS (mean MFI, x10^3)
c
RV-C15 infection (AUC)
HS (mean MFI, x10^3)
mono17
mono24
mono50
mono29
mono25
mono53
mono67
RV-C15
HS Expression
d
% Infection
RV-C11
CDHR3
CDHR3 (MFI)
HS
HS (MFI, x10^3)
mono25 mono50 mono24 mono67 mono17 mono29 mono11 mono53 poly mono56 mono39 mono9
e
RV-C11 Infection (mean %)
Spearman ρ = 0.81, p = 0.0016
CDHR3 (mean MFI)
Spearman ρ = -0.47, p = 0.13
HS (mean MFI, x10^3)
f
% Infection (C15)
mono25
AUC: 34.8
mono53
AUC: 30.2
mono67
AUC: 14.1
Log10 Virus Dilution
g
% Infection (C15)
R² = 0.07, p = 7.37E-56
Passage# (mono29)

**Figure S3. Detailed characterization of CDHR3$^{C529Y}$-expressing HeLa monoclonal clones.** (a) Flow cytometry analysis of heparan sulfate (HS) expression in RV-C15 permissive and non-permissive clones. Infection efficiency is shown as green circles (% mGL-positive cells), and HS expression as red triangles (mean fluorescence intensity; MFI). Each point represents the mean of two technical replicates from an independent experiment. Colored bars indicate the overall mean from all experiments (n=1 – 3). (b) Correlation between RV-C15 infection permissiveness (% mGL-positive cells) and CDHR3 expression (MFI) (left) or HS expression (right). Mean values from three independent experiments are shown. Clones with infection above 30% are highlighted in green. Correlation was assessed by Spearman's rank test. A loess regression line is shown for visualization only. (c) Correlation between RV-C15 infection permissiveness (AUC) and HS expression (MFI) across monoclonal clones. (d) Flow cytometry analysis of surface CDHR3 expression (left) or HS expression (right) in RV-C11 permissive and non-permissive clones. Infection efficiency is shown as green circles (% mGL-positive cells), and CDHR or HS expression as red triangles (mean fluorescence intensity; MFI). Each point represents the mean of two technical replicates from an independent experiment. Colored bars indicate the overall mean from all experiments (n=1 – 2). (e) Correlation between RV-C11 infection permissiveness (% mGL-positive cells) and CDHR3 expression (MFI) (left) or HS expression (right). Mean values from three independent experiments are shown. Clones with infection above 15% are highlighted in green. Correlation was assessed by Spearman's rank test. A loess regression line is shown for visualization only. (f) Related to Fig. 1d. RV-C15 infection profiles of permissive clones in dose-response assays. Infection is quantified as area under the curve (AUC). (g) Long-term monitoring of mono29 showing sustained permissiveness to RV-C15. Mean infection % from each passage was plotted against passage number, and linear regression was applied. $R^2$ and p values are shown (R2=0.07, p=7.37E-56, total n=3384). The dash line represents linear regression fit.

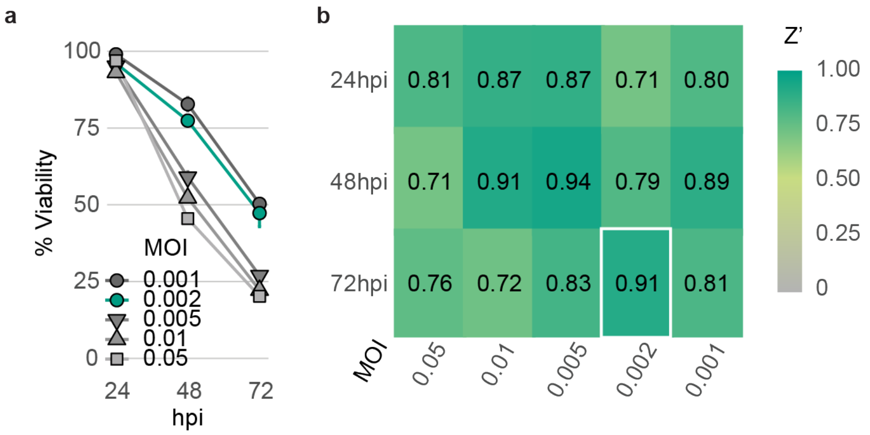


c

| Virus | Compound | Mean IC50 ± SD (µM) | Inter-operator CV (%) | ANOVA p-value | Z′ (mean ± SD) | N (plates) |
|---|---|---|---|---|---|---|
| RV-A16 | BF738735 | 0.017 ± 0.001 | 8.1% | 0.40600 | 0.88 ± 0.06 | 361 |
| RV-A16 | Pirodavir | 0.013 ± 0.001 | 10.1% | 0.18200 | | |
| RV-A16 | Rupintrivir | 0.004 ± 0.000 | 2.5% | 0.65400 | | |
| RV-B14 | BF738735 | 0.051 ± 0.004 | 8.6% | 0.70100 | 0.89 ± 0.05 | 363 |
| RV-B14 | Pirodavir | 0.028 ± 0.005 | 19.7% | 0.38000 | | |
| RV-B14 | Rupintrivir | 0.005 ± 0.001 | 15.2% | 0.16900 | | |
| RV-C15 | BF738735 | 0.029 ± 0.007 | 25.3% | 0.00873 | 0.80 ± 0.10 | 363 |
| RV-C15 | Rupintrivir | 0.014 ± 0.002 | 12.8% | 0.12900 | | |

d

| Virus | Compound | Operator | Mean IC50 ± SD (µM) |
|---|---|---|---|
| RV-A16 | BF738735 | Op1 | 0.018 ± 0.006 |
| RV-A16 | BF738735 | Op2 | 0.016 ± 0.004 |
| RV-A16 | Pirodavir | Op1 | 0.012 ± 0.003 |
| RV-A16 | Pirodavir | Op2 | 0.014 ± 0.003 |
| RV-A16 | Rupintrivir | Op1 | 0.004 ± 0.001 |
| RV-A16 | Rupintrivir | Op2 | 0.004 ± 0.000 |
| RV-B14 | BF738735 | Op1 | 0.048 ± 0.033 |
| RV-B14 | BF738735 | Op2 | 0.054 ± 0.027 |
| RV-B14 | Pirodavir | Op1 | 0.024 ± 0.016 |
| RV-B14 | Pirodavir | Op2 | 0.032 ± 0.016 |
| RV-B14 | Rupintrivir | Op1 | 0.005 ± 0.001 |
| RV-B14 | Rupintrivir | Op2 | 0.006 ± 0.002 |
| RV-C15 | BF738735 | Op1 | 0.024 ± 0.010 |
| RV-C15 | BF738735 | Op2 | 0.034 ± 0.007 |
| RV-C15 | Rupintrivir | Op1 | 0.012 ± 0.004 |
| RV-C15 | Rupintrivir | Op2 | 0.015 ± 0.004 |

**Figure S4. Optimization of antiviral assay in 96-well format.** (a) Related to Fig. S2a. Infection kinetics and cytopathic effects of RV-C15a-mGL in HeLa-CDHR3$^{C529Y}$ mono29 cells across multiple MOIs and time points. Data are presented as mean ± SD (n=3). (b) Heatmap of Z' factors across different MOIs and time points. The selected condition for subsequent antiviral assay is outlined in white. (c) Inter-operator reproducibility of assay performance across >360 plates. Z′ factors and $IC_{50}$ values of reference compounds (BF738735, Pirodavir, Rupintrivir) were compared between two independent operators. (d) Intra-operator reproducibility of assay performance.

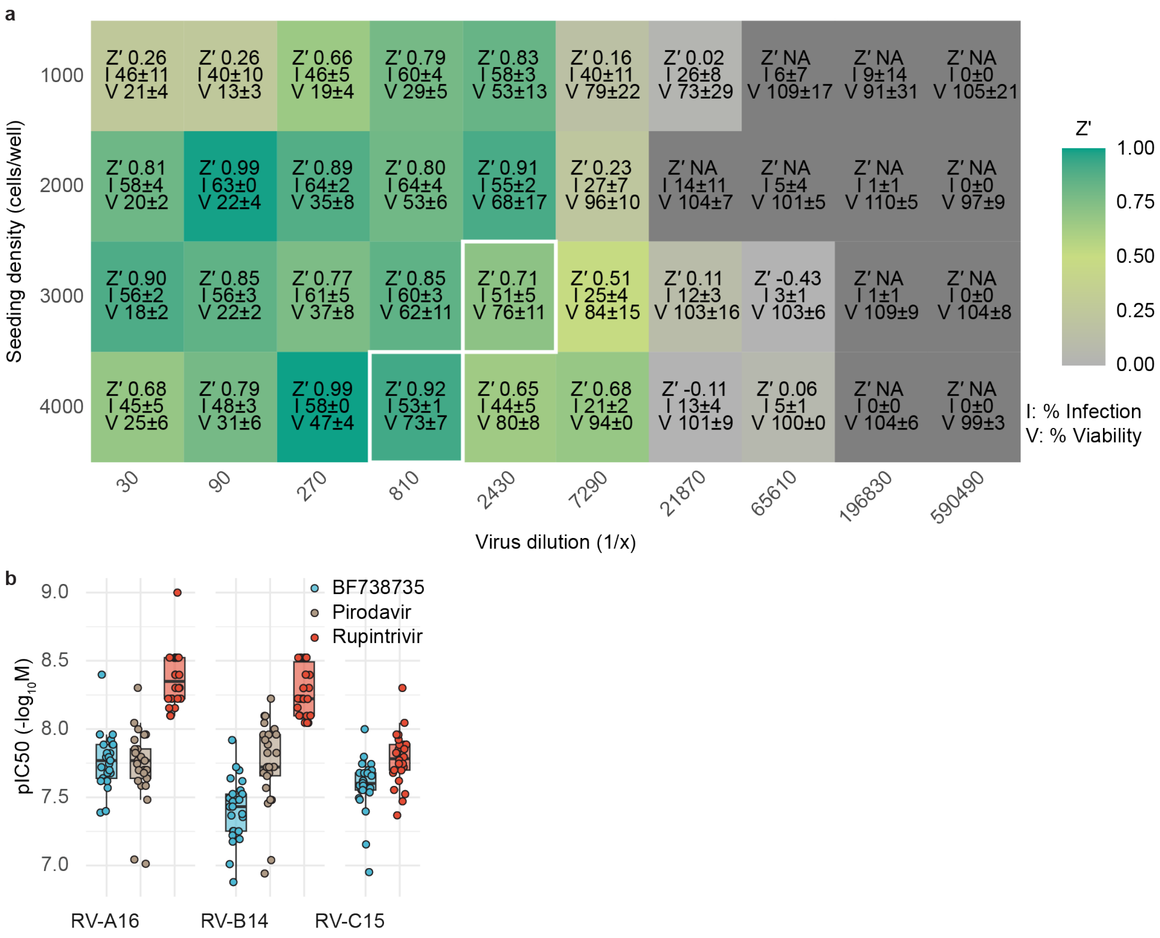


**Figure S5. Miniaturization of the assay to 384-well format.** (a) Heat map of Z' factors across different seeding densities (1000 – 4000 cells/well) and viral dilutions. Values represent mean ± SD of three independent replicates. Two optimal conditions meeting all three criteria (Z' > 0.7, infection > 50%, and viability > 70%) are indicated in white borders. (b) Related to Fig. 3c. Distribution of $pIC_{50}$ values ($-\log_{10} IC_{50}$, M) for BF738735, pirodavir, and rupintrivir across >100 384-well RV-A16, RV-B14, and RV-C15 assays. Each dot represents an individual run; boxes indicate median, interquartile range (IQR), and whiskers (1.5× IQR).

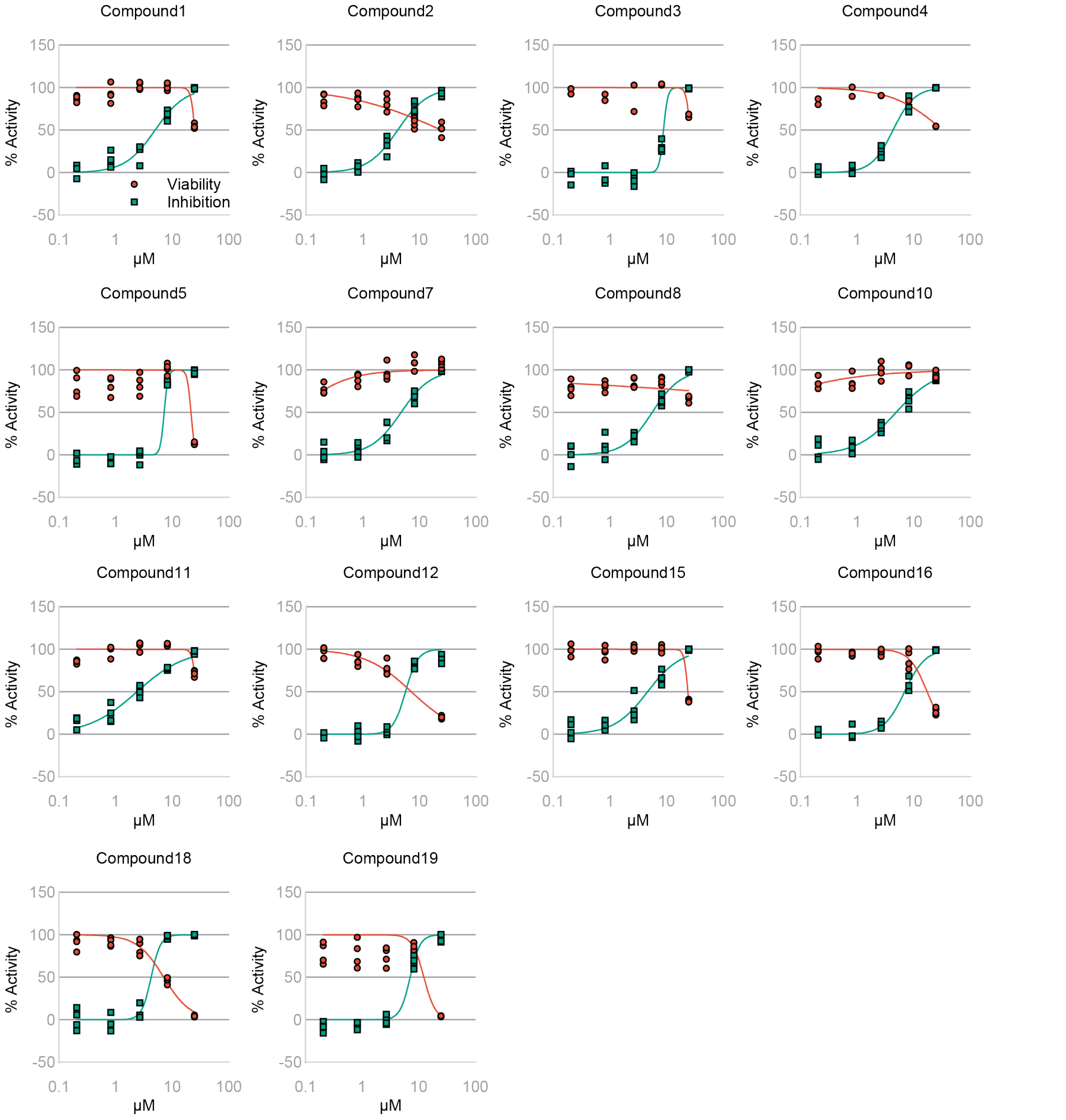


**Figure S6. Validation of primary screening hits.** Related to Fig. 4d. IC50 and CC50 curves from mini dose-response assays (0.2 – 25 μM, four replicates) of 14 out of 19 primary hits identified in the pilot screen.

**Tables**

**Table S1.** Antiviral activity of 19 primary hits confirmed in RV-C15a-mGL mini dose–response assays.

| **CompoundID** | **EC50 (μM)** | **CC50 (μM)** | **SI** |
|---|---:|---:|---:|
| Compound1 | 4.63 | 24.90 | 5.38 |
| Compound2 | 10.14 | 24.90 | 2.46 |
| Compound3 | 4.07 | 24.83 | 6.10 |
| Compound4 | 5.54 | 15.91 | 2.87 |
| Compound5 | 4.30 | 24.90 | 5.79 |
| **Compound6** | **3.31** | **24.90** | **7.51** |
| Compound7 | 4.67 | 24.90 | 5.33 |
| Compound8 | 4.80 | 24.90 | 5.19 |
| **Compound9** | **2.35** | **24.90** | **10.62** |
| Compound10 | 5.59 | 24.90 | 4.46 |
| Compound11 | 5.12 | 18.02 | 3.52 |
| Compound12 | 5.64 | 7.19 | 1.28 |
| **Compound13** | **1.73** | **24.90** | **14.42** |
| **Compound14** | **3.13** | **24.90** | **7.94** |
| Compound15 | 4.67 | 22.42 | 4.80 |
| Compound16 | 6.90 | 17.49 | 2.54 |
| **Compound17** | **2.98** | **24.90** | **8.37** |
| Compound18 | 4.42 | 7.20 | 1.63 |
| Compound19 | 6.84 | 12.57 | 1.84 |

**Table S2.** Primers and probe used in this study.

| Purpose | Primer/Probe | Sequence (5' – 3') |
|---|---|---|
| Cloning pLenti-CDHR3$^{C529Y}$-IRES-Blast | CDHR3 F | ACACAGGTGTCGTGACGCGGATCCATGCAGGAAGCAATCATTC |
| | CDHR3 R | TCACTTTCCTGGGTGTGGTTT |
| | pLenti F | CCCCAAACCACACCCAGGAAAGTGATGCATGCCCCTCTCC |
| | pLenti R | GGATCCCGCGTCACGACA |
| Cloning pRV-C15 | pUC19 470F | AGCTGTTTCCTGTGTGAAATTG |
| | pUC19 2676R | CCTCGTGATACGCCTATTTTTATA |
| Cloning pRV-C15a | C15-3A F | AAGAGGGATTTTAATTATGTGCATTA |
| | C15-VP1 R | TTGTTGTTGGTTACTATGGTAAC |
| | C15-VP1 F | TACCATAGTAACCAACAACAAAGGGTTGATGCAAATAATGTTC |
| | C15-3A R | AATGCACATAATTAAAATCCCTCTTTATGGTTGAGTTTGCTTTACCTA |
| Cloning pRV-C15a-mGL | C15-mGL-2A F | CTCATCAGTTCTGCGGGACCGAGCGACATG |
| | C15-mGL-VP1 R | ACTAGGTCCAGCTGAGCTGATGAGGGGTCTGG |
| | mGL F | CCTCATCAGCTCAGCTGGACCTAGTGTGAGCAAGGGCGAGGAG |
| | mGL R | CGCTCGGTCCCGCAGAACTGATGAGCTTGTACAGCTCGTCCATGTC |
| Cloning pUC19-CMV/T7 | pUC19 427F | TAGAGGATCCCCGGGTACC |
| | pUC19 426R | GAGTCGACCTGCAGGCAT |
| | CMV F | AGCTTGCATGCCTGCAGGTCGACTCGACATTGATTATTGACTAGTTATTAATA |
| | T7-HHR R | TCATCAGTTAAATCTCCCTATAGTGAGTCGT |
| | HDR F | CAACATTCCGAGGGGACCGTCCCCTCGGTAATGGCGAATGGG |
| | SV40pA R | GAGCTCGGTACCCGGGGATCCTCTAACATGATAAGATACATTGATG |
| Cloning pRV-C11a-mGL | UC-1 (2) | GGCCGGCATGGTCCCAGCCTCCTCGCTGGCGCCGGCTGGGCAACATT |
| | UC-2 (2) | GACGGAGTCTAGACTCCGTT |
| Cloning pRV-C41a-mGL | UC-1 | GGCCGGCATGGTCCCAGCCTCCTCGCTGGCGCCGGCTGGGCAACATT |
| | UC-2 C41 | CCCAGTTTTAAGACGGAGTCTAGACTCCGTT |
| RT-qPCR (3) | Panenterhino F | AGCCTGCGTGGCKGCC |
| | Panenterhino R | GAAACACGGACACCCAAAGTAGT |
| | Panenterhino Probe | /56-FAM/CTCCGGCCC/ZEN/CTGAATGYGGCTAA/3IABkFQ/ |
| RT-PCR RV-C15a-mGL | C15-VP1 F | GGACTTTGGACAATAGTGGTG |
| | C15-2A R | GAGTAGTAGCAACCAGCAGTG |

**Table S3.** Synthesized DNA fragments used in this study.

| Fragment | Description | Sequences (5' – 3') |
|---|---|---|
| C15 gBlocks1 | pUC19 overhang-T7 promoter-5'UTR to VP3 | CTATAAAAATAGGCGTATCACGAGGCAGCTTGCATGCCTGCAGGTCGACTCTAGAGGATCTCGATCCCGCGAAATTAATACGACTCACTATAGGTTAAAACTGGGTATAGGTTG……ATGGGACTTGGGCCTGCAGTCGTCCTGCACCATTGTGGTACCATGGATTT |
| C15 gBlocks2 | VP3 to 2C | TGCACCATTGTGGTACCATGGATTTCATCTGGTTTCTACCGCCGCACCAA……GTCACCACCAACAGTCACAATACCTGAGGCTATAAAACGGCGATTCTTTC |
| C15 gBlocks3 | 2C to 3D | GAGGCTATAAAACGGCGATTCTTTCTGGATGCCGATCTCATCACAACATC……TCTCTACACCCCTCCTTACTCATTGTTGTACAGGCAATGGATAGATCAAT |
| C15 Ultramer1 | 3D-PolyA(50)-pUC19 overhang | TTGTACAGGCAATGGATAGATCAATTCATATAAACAAATAATATAATACAATGTTTGAGTAGAACTTAGTATTATAAAAAAAAAAAAAAAAAAAAAAAAAAAAAAAAAAAAAAAAAAAAAAAAAACCGCGGCCTGGATCCCCGGGTACCGAGCTCGAATTCGTAATCATGGTCATAGCTGTTTCCTGTGTGAAATTGTTA |
| pUC19-CMV/T7 Ultramer 1 | | CACTATAGGGAGATTTAACTGATGAGTCCGTGAGGACGAAACGGAGTCTAGACTCCGTCGCTACCATCTAACCGGTTCATTCTCAAAAAAAAAAAAAAAAAAAAAAAAAAAAAAAAAAAAAAAAAAAAAAAAAAGGCCGGCATGGTCCCAGCCTCCTCGCTGGCGCCGGCTGGGCAACATTCCGAGGGGACCGTCCCCT |
| pUC19-CMV/T7 Ultramer 2 | | CGGTAATGGCGAATGGGACCTAGCATAACCCCTTGGGGCCTCTAAACGGGTCTTGAGGGGTTTTTTGAACTTGTTTATTGCAGCTTATAATGGTTACAAATAAAGCAATAGCATCACAAATTTCACAAATAAAGCATTTTTTTCACTGCATTCTAGTTGTGGTTTGTCCAAACTCATCAATGTATCTTATCATGTTAGAG |
| C11 gBlocks1 | pUC19-CMV/T7 overhang-5'UTR to VP2 | GACGAAACGGAGTCTAGACTCCGTCTTAAAACTGGATACAGGTTGTTCCC……TGGTTTACTACTTGTTGCTATAGTACCAGAGCACCAGTTAGCATACGTCG |
| C11 gBlocks2 | VP2 to VP1 | CCAGAGCACCAGTTAGCATACGTCGGTGGCACATATGCTAATGTTGGGTA……ACCAATACACACTACCAAACCAACTGCTCTATCGGCAATGGAGATTGGAG |
| C11 gBlocks3 | VP1-mGL-2C | GCTCTATCGGCAATGGAGATTGGAGCATCTTCTGATGTAAAACCAGAAGA……GGAAGAGCTTTACCAGTGACTTTGTGTTGGCTAGTACAAATTTAAACCAG |
| C11 gBlocks4 | 2C to 3D | GTTGGCTAGTACAAATTTAAACCAGTTAAGCCCCCCCACTGTAACAATAC……AACAGTATGATGACTTTGTCCAGAAAGTTCGATCCGTCCCTGTAGGTCGA |
| C11 Ultramer1 | 3D-PolyA(50)-pUC19-CMV/T7 overhang | AGTTCGATCCGTCCCTGTAGGTCGAACCTTGTACTTACCACCATATGAGTACTTAGAAAGAAGATGGATAGATAAATTCATATAATTAGATATGGTATTCAAGTTGTTTAGAATTGACTATCATAAAAAAAAAAAAAAAAAAAAAAAAAAAAAAAAAAAAAAAAAAAAAAAAAAGGCCGGCATGGTCCCAGCCTCCTCG |
| C41 gBlocks1 | pUC19-CMV/T7 overhang - 5'UTR to VP3 | GTCTAGACTCCGTCTTAAAACTGGGTAGGGGTTGTTCCCACCCCTACCAC……TAGCGCCTGCCCTGACATGTCTGTTAGGATGATGAGGGATAGTCCAATGA |
| C41 gBlocks2 | VP3-mGL-2C | AGGATGATGAGGGATAGTCCAATGATGAAGCAAGAAGGGAAGCTCCAAAA……TTACTACACGGGAACCCCGGTTGTGGTAAATCTCTTGCTACGTCCGTCAT |

| | | |
|---|---|---|
| C41 gBlocks3 | 2C to 3D | GTAAATCTCTTGCTACGTCCGTCATTGCACGTGGCCTCACAACAGAAGCA……TCTATGAAGAATTCAGATCTAAGATCAGAAGTACAGCAGTAGGAAGATCC |
| C41 Ultramer1 | 3D-PolyA(50)-pUC19-CMV/T7 overhang | CAGAAGTACAGCAGTAGGAAGATCCTTATACACCCCCCCTTACTCTGTACTGTATAGACAGTGGATTGATTTATTTGTTTAGATAATTAAATATAATACAGTGTTTGCTTAGAACTTAGTATTATAAAAAAAAAAAAAAAAAAAAAAAAAAAAAAAAAAAAAAAAAAAAAAAAAAGGCCGGCATGGTCCCAGCCTCCTCG |

Green: T7 promoter
Red: overlapping sequences
Blue: XmaI site
Underline: Virus 5'UTR or 3'UTR

**SI References**


1. Y. A. Bochkov, *et al.*, Cadherin-related family member 3, a childhood asthma susceptibility gene, mediates rhinovirus C binding and replication. *Proc Natl Acad Sci U A* **112**, 5485–5490 (2015).

2. W.-S. Choi, *et al.*, Development of a Universal Cloning System for Reverse Genetics of Human Enteroviruses. *Microbiol. Spectr.* **11**, e0316722 (2023).

3. C. Tapparel, *et al.*, New Respiratory Enterovirus and Recombinant Rhinoviruses among Circulating Picornaviruses. *Emerg. Infect. Dis.* **15**, 719–726 (2009).
